# XMRF: An R package to Fit Markov Networks to High-Throughput Genetics Data

**DOI:** 10.1101/032219

**Authors:** Ying-Wooi Wan, Genevera I. Allen, Yulia Baker, Eunho Yang, Pradeep Ravikumar, Zhandong Liu

## Abstract

**Motivation:** Technological advances in medicine have led to a rapid proliferation of high-throughput “omics” data. Tools to mine this data and discover disrupted disease networks are needed as they hold the key to understanding complicated interactions between genes, mutations and aberrations, and epi-genetic markers.

**Results:** We developed an R software package, `XMRF`, that can be used to fit Markov Networks to various types of high-throughput genomics data. Encoding the models and estimation techniques of the recently proposed exponential family Markov Random Fields (Yang et al., 2012), our software can be used to learn genetic networks from RNA-sequencing data (counts via Poisson graphical models), mutation and copy number variation data (categorical via Ising models), and methylation data (continuous via Gaussian graphical models).

**Availability:** XMRF is available from the CRAN Project and Github at: https://github.com/zhandong/XMRF

## Introduction

Markov random fields (MRFs) are a popular tool for estimating relationships between genes, finding regulatory pathways, and visually depicting genetic networks. Estimating sparse, high-dimensional undirected graphical models, or Markov Networks, linking a set of *p* genes measured on *n* samples has been well studied for the Gaussian graphical model (GGM) [1] and also the binary or categorical Ising model [2]. As many high-throughput genomic data sets such as counts observed in Next Generation Sequencing data, are not approximately Gaussian or binary, existing methods for graphical models are greatly limited. To address this, a recent line of work has proposed a parametric family of graphical models based on univariate exponential family distributions with a rich theoretical foundation [3, 4]. These models can be used to estimate genetic networks from a variety of data types: gene expression networks based on RNA Sequencing via Poisson-family graphical models, mutation and aberration networks via Ising graphical models, and epi-genetic networks via Gaussian graphical models. In this paper, we introduce an R software package, `XMRF`, that encodes models and estimation techniques for fitting exponential family Markov Networks to high-throughput genomics data as well as software to pre-process genetic data and visualize the resulting genetic networks.

## Algorithms and Implementations

Recently, we proposed a novel class of MRF models [3] constructed by assuming that all node-conditional distributions are univariate exponential families. These then yield a class of models appropriate for a variety of data types such as counts, categorical, continuous, and skewed continuous variables found in high-throughput genomics data. Further, we proposed a simple way of learning the network structure of these exponential family MRFs by maximizing the penalized node-conditional likelihoods; these methods come with strong theoretical guarantees for sparse graph estimation as discussed in [3]. As the models are constructed via exponential families, the so-called neighborhood selection problems for each node reduce to that of an *ℓ*_1_ penalized and possibly constrained generalized linear model (GLM) [3].

Suppose *X* = (*X*_1_,…,*X_p_*) is a random vector, with each variable *X_i_* taking values in a set χ. Suppose *G =* (*V, E*) is an undirected graph over *p* nodes corresponding to the *p* variables; the corresponding graphical model is a set of distributions that satisfy Markov independence assumptions with respect to the graph. Then, conditioned on all other variables, *X_s_* is distributed according to an exponential family distribution [4]: 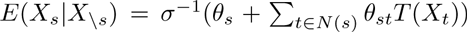, where *θ_st_* is the weight parameter denoting an edge between *X_s_* and *X_t_*, *T*(·) is the sufficient statistics function, and *N*(*s*) is the neighborhood, or set of edges extending out from, *X_s_*. This gives the following conditional density: 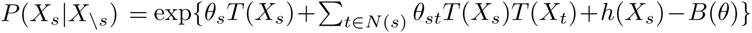. Based on the Hammersley-Clifford theorem, these conditional densities give the following joint density, or MRF, over the set of nodes to form our *Exponential Family Graphical Model:*

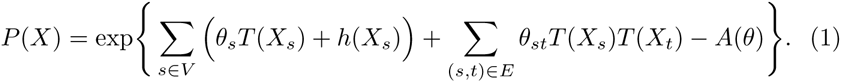

Here, *A*(*θ*) is a log-normalization term ensuring that *P*(*X*) is a proper density. We then fit this model using penalized conditional maximum likelihood estimation which corresponds to a neighborhood selection problem for the *s^th^* node of the following form:

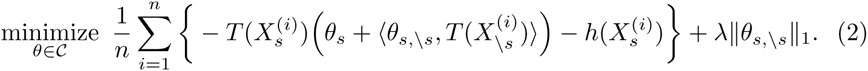

Here, *C* is the constraint region of the parameter space as discussed further for specific examples in [4]. Notice that this cooresponds to fitting a penalized GLM.

In the `XMRF` package, we implement the neighborhood selection graph estimation technique by proximal or projected gradient descent using warm-starts over the range of regularization parameters, *λ*. Note also that each node-neighborhood problem is completely independent and can hence be computed in parallel; this is achieved using the default parallel support and the `snowfall` package [5] in R. The max-rule is used to construct the network from all neighborhood estimates. Selecting the sparsity of the network corresponds to selecting λ. We implement two data-driven approaches to do so: StARS which is computed over a range of λ values [6], and a stability-based approach for a single value of λ which for every edge, computes the proportion of times the edge is selected in the model (termed the stability score) over many bootstrap re-samples [7]. Our package also implements Gibbs samplers to simulate data for our exponential family MRF models. Finally, our package includes functions for pre-processing sequencing data [8] and includes a host of network visualization and network manipulation strategies through a dependency on the `igraph` package [9]. Our package is developed under statistical computing environment R and compatible to be executed in version 3.0.2 or later.

## Results and Discussion

To estimate the network structures from different types of high-throughput genomics data, our package consists of one main function, the `XMRF()` function, for which many families of distributions are possible, `XMRF(…, method=“GGM”)`: the `GGM` for family of Gaussian graphical models, the `ISM` for Ising models, `PGM`, `TPGM`, `SPGM`, `LPGM` for Poisson families of models as described in [8, 10]. For genomic networks based on sequencing data, we recommend using the `LPGM` variant, but all methods are described in the package vignette. Table 1 summarizes each of the main families in our `XMRF()` function as well as our recommendation for which family to use for various types of high-throughput genomics data.

**Table 1.** Recommended families to use in our XMRF package.

| Genetics Data | Type | XMRF Family |
| --- | --- | --- |
| RNA-Seq or miRNA-Seq | Counts | LPGM or SPGM |
| Microarray or Methylation | Continuous | GGM |
| Mutations or CNVs | Binary | ISM |

The `XMRF()` function will return an object of `GMS` class representing the fitted models. The `GMS` object contains the list of fitted networks, the stability of each fitted network, the full regularization path, and the index of the optimal network. The default `plot` method of `GMS` class enables drawing the learned network in graphical format and saves the output to a PDF document. The package also includes `plotGML` function to write the learned network in graph modeling language, which can be imported to Cytoscape [11] for further visualization customization.

### Choosing the Right Graphical Model for Genomics Data

The development of high-throughput technology, such as microarray, SNP array, array-CGH, methylation array, exome-sequencing, and RNA-sequencing, has generated a wide variety of genetics data. Each of these genetics data varies in data types. For example, next generation sequencing (RNA-Seq and miRNA-Seq) data are read counts; expression profiles from microarray and methylation array are continuous values; mutations and CNVs are usually represented in binary, with value one represents the gene is mutated, amplified or deleted in the patient, or value zero otherwise (Table 1). To accurately estimate the underlying network structure from these data types, one needs to apply the right network inference algorithm based on the platform-specific data distribution.

In `XMRF`, data are modeled using their native distribution instead of normalizing the data to follow a Gaussian distribution. To accomplish this, our package implements methods for three families: Gaussian graphical models (`GGM`), Ising models (`ISM`), and Poisson family graphical models including regular Poisson graphical models (`PGM`) as well as several variants of the Poisson family of models such as the truncated Poisson (`TPGM`), sub-linear Poisson (`SPGM`), and local Poisson (`LPGM`) [8]. Note that [10] proposed all these variants of the Poisson family as the regular Poisson graphical model only permits negative conditional dependencies between nodes; each of these variants relaxes restrictions resulting in both positive and negative conditional dependencies. For genomic networks based on sequencing data, we recommend using the `LPGM` variant as proposed in [8], noting that this local model closely approximates the proper MRF distribution of the `SPGM` formulation [10].

### Poisson Graphical Model for NGS data

Count data generated by next generation sequencing (NGS) is a good example of why parametric families of Markov Networks beyond Gaussian graphical models are needed. This read count data is highly skewed and has large spikes at zero so that standardization to a Gaussian distribution is impossible. Here, we demonstrate through a real example how to process NGS data so that we can use Poisson family graphical models to learn the network structure. Our processing pipeline is given in [12] and encoded in the `processSeq` function of the package.

The level 3 RNA-Seq (UNC Illumina HiSeq RNASeqV2) data consisting of 445 breast invasive carcinoma (BRCA) patients from the Cancer Genome Atlas (TCGA) project [13] was obtained. The set of 353 genes with somatic mutations listed in the Catalogue of Somatic Mutations in Cancer (COSMIC) were further extracted [14]. The data was prepared and stored as the `brcadata` data object included in the package. The values in this data object are the normalized read counts (RSEM) obtained from TCGA data download portal, representing the mRNA expression profiles of the genes. The data matrix is of dimension *pxn*. Figure 1 shows that the data is more appropriate for Poisson family graphical models after being preprocessed with the `processSeq` function as given in the following code snippets:

~~~
> library(XMRF)
> data(’brcadata’)
> brca = t(processSeq(t(brcadat), PercentGenes=1))
~~~

**Figure 1.**
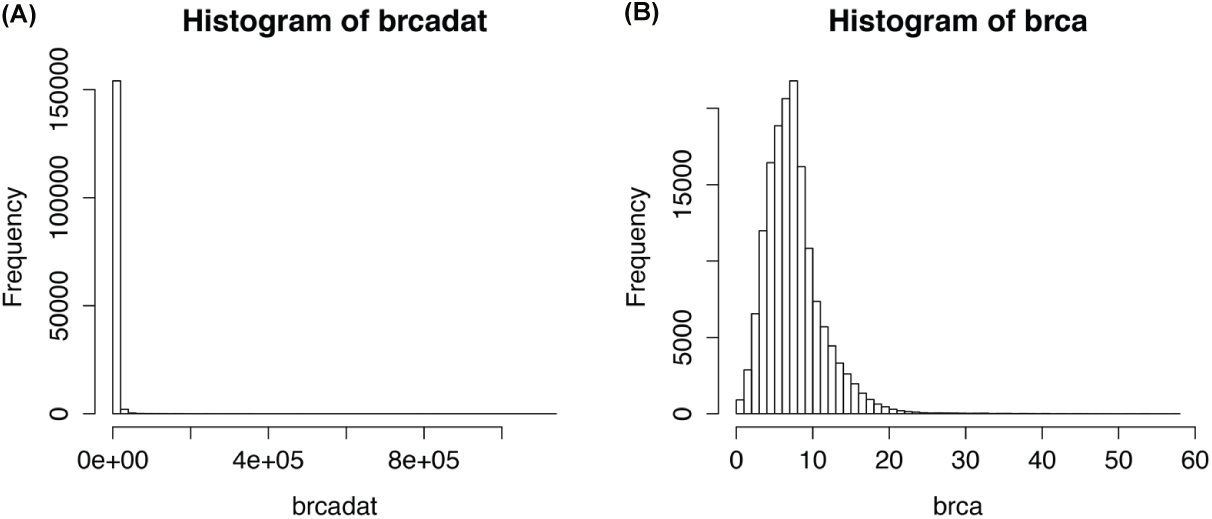
Distribution of TCGA BRCA RNA-Seq data before (A) and after (B) preprocessing. The latter gives a distribution more appropriate for Poisson family graphical models.

To estimate the underlying network structure of the count-valued data, `XMRF` implements four different models from the Poisson family graphical models: regular Poisson graphical model (`PGM`) that only permits negative conditional dependencies, truncated Poisson (`TPGM`), sub-linear Poisson (`SPGM`), and local Poisson (`LPGM`) [8]. The latter three models are variants of the Poisson family that relax restrictions as imposed in a regular Poisson model, resulting in both positive and negative conditional dependencies [10]. `TPGM` should be used if one wants to truncate the large counts observed in NGS dataset. Alternatively, `SPGM` implements a sub-linear truncation for the NGS data which gives a softer reduction on large counts. `LPGM` is a faster algorithm that approximates the Markov Network while preserving both positive and negative relationship [8].

In practice, we choose `LPGM` since it is the fastest and most flexible way to capture both positive and negative dependencies [12]. As an example, we applied `XMRF` functions to study the relationships between 353 genes with somatic mutations cataloged in the COSMIC cancer gene census database. Gene expression data measured via RNA-Seq for 445 samples was acquired from the Cancer Genome Atlas (TCGA). The `processSeq` function was used to process the sequencing data so that our Poisson graphical models are appropriate [8]. The estimated network, Figure 2, includes multiple associations reported in published literature, such as the associations of FOXA1, CCND1, and PBX1 with GATA3, link between ERBB2 and CDK12, and others. These results validate the utility of our methods and algorithms implemented in the package for finding gene interactions.

~~~
> library(XMRF)
> data(’brcadat’)
> brca = t(processSeq(t(brcadat), PercentGenes=1))
> lambda = 0.1 * lambdaMax(t(as.matrix(brca)))*
  sqrt(log(nrow(brca))/ncol(brca))
> brca.lpgm <- XMRF(brca, method=“LPGM”, lambda.path=lambda,
 th=0.005, sth=0.9)
> plotGML(brca.lpgm, fn=“brcanet.gml”, weight=TRUE, vars=rownames(brca))
~~~

**Figure 2.**
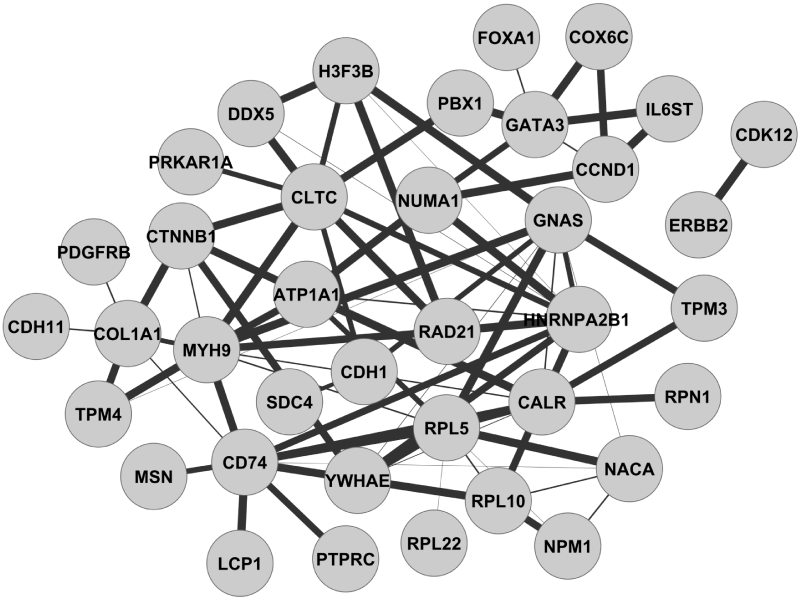
Inferred relationships between cancer census genes from TCGA breast cancer patients. The width of edges reflects the strength of inferred relationships.

### Gaussian Graphical Model for Expression Arrays

When genomics data is profiled with microarrays, such as with mRNA arrays, miRNA arrays, or methylation arrays, Gaussian graphical models should be used to estimate the network structures. Similar workflows as presented in last section can be applied to fitting Gaussian Markov Networks to data that approximately follows a multivariate Gaussian distribution.

Here, we give an example of the work-flow of learning gene networks associated with kidney renal clear cell carcinoma (KIRC) from tumor patients [15]:

1. Obtain gene expression data for KIRC, profiled with mRNA microarray.
2. Obtain data for only tumor samples.
3. Filter genes so that the top 5% of variable genes remain.
4. Use the `XMRF` function to learn the network structure. Note that it is always good practice to visualize the data to confirm the distributional family before model fitting. In this example as shown in Figure 3, the data follows a Gaussian distribution and thus fitting a Gaussian Graphical model is appropriate.
5. Write the network in GML format and view the network via Cytoscape (Figure 4).

Code snippets for the above work-flow are provided as follows:

~~~
> # 1. Get TCGA data
> # Obtain mRNA array gene expression data for KIRC patients
> library(TCGA2STAT)
> kirc<- getTCGA(disease=“KIRC”, data.type=“mRNA_Array”)
# 2. Obtain data for tumor samples
> kirc.tum <- SampleSplit(kirc$dat)$primary.tumor
> # 3. Filter genes to remain those of top 5% most varied genes
> var <- apply(kirc.tum, 1, var)
> nac <- apply(kirc.tum, 1, function(x) sum(is.na(x)))
> kirc.tum.gd <- kirc.tum[var >= quantile(var, probs=0.95, na.rm=T)
                        & !is.na(var) & nac==0,]
> # Take a look at the data to confirm distribution family
> hist(kirc.tum.gd, breaks=20)
~~~

**Figure 3.**
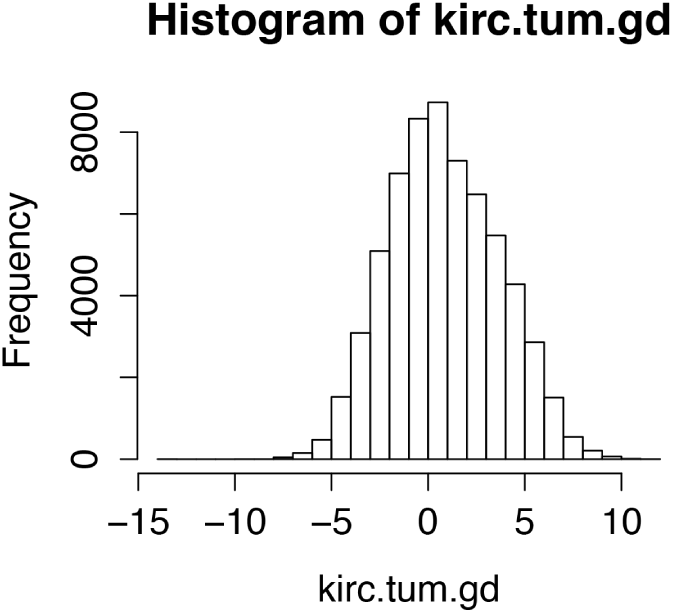
Distribution of mRNA expression profiled with micrarray from KIRC tumor samples.

~~~
> # 4. Fit the data to Gaussian graphical model
> kirc.tum.fit <- XMRF(kirc.tum.gd, method=“GGM”, N=100,
                                    stability=“STAR”, nlams=10, beta=0.001)
> # Visualized the gene network
> plotGML(kirc.tum.fit, fn=“kirc.tum.array.gml”, i=2,
                              weight=TRUE, vars=rownames(kirc.tum.gd))
~~~

**Figure 4.**
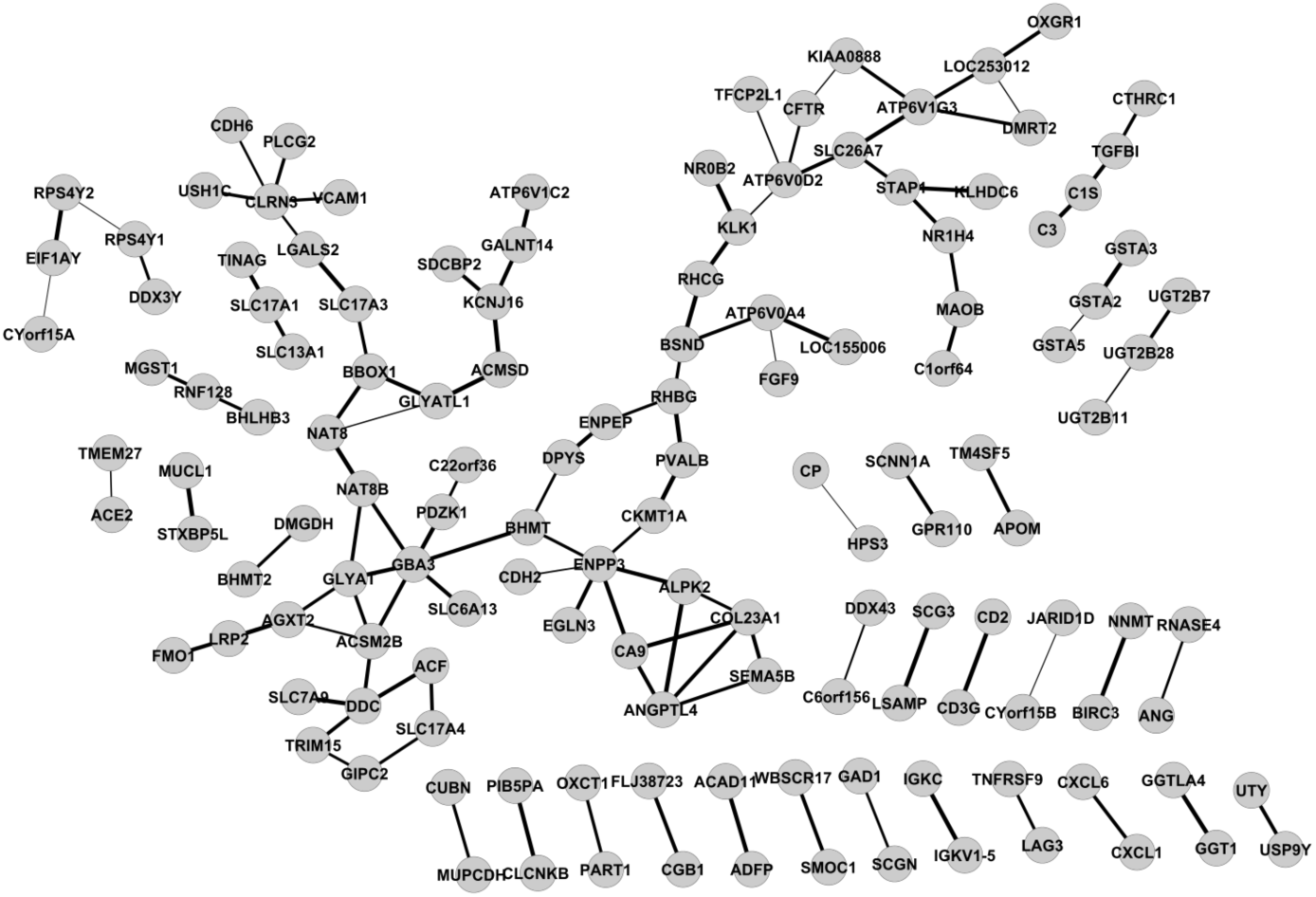
KIRC expressed gene networks estimated by GGM via `XMRF(…,method=“GGM”)` for mRNA expression data.

### Ising Graphical Model for Mutation Data

To fit Markov Networks to binary data, the `XMRF` function with `method=“ISM”` can be used. In this section, we give an example of fitting an Ising model to simulated data with a lattice graph as well as estimating interactions among mutated genes in TCGA lung squamous cell carcinoma (LUSC) samples [16].

#### Learning a Lattice Graph from Simulated Data

In the following example, a multivariate binary data matrix of 400 observations that will give a 5 x 5 grid graph will be simulated. Our `XMRF` Ising model is fit to infer the lattice network from simulated data. Models were fitted over a path of 20 regularization values. StARS stability selection with 100 iterations were used to select the optimal network. Figure 5 shows that the estimated optimal network shown (Figure 5(B) and (D)) is almost identical to the true simulated network (Figure 5(A) and (C)). This result shows that the Ising graphical model implemented in our package could correctly identify the relationships between variables from binary data.

~~~
> n = 400
> p = 25
> simdat <- XMRF.Sim(n=n, p=p, model=“ISM”, graph.type=“lattice”)
> ismfit <- XMRF(simdat$X, method=“ISM”, N=100, nlams=20,
                 stability=“STAR”, th=0.1, beta=0.1)
> par(mfrow=c(2,2))
> image(simdat$B)
> image(ismfit$network[[ismfit$opt.index]])
> ml = plotNet(simdat$B, fn=““)
> ml = plot(ismfit, fn=“”, mylayout=ml)
~~~

**Figure 5.**
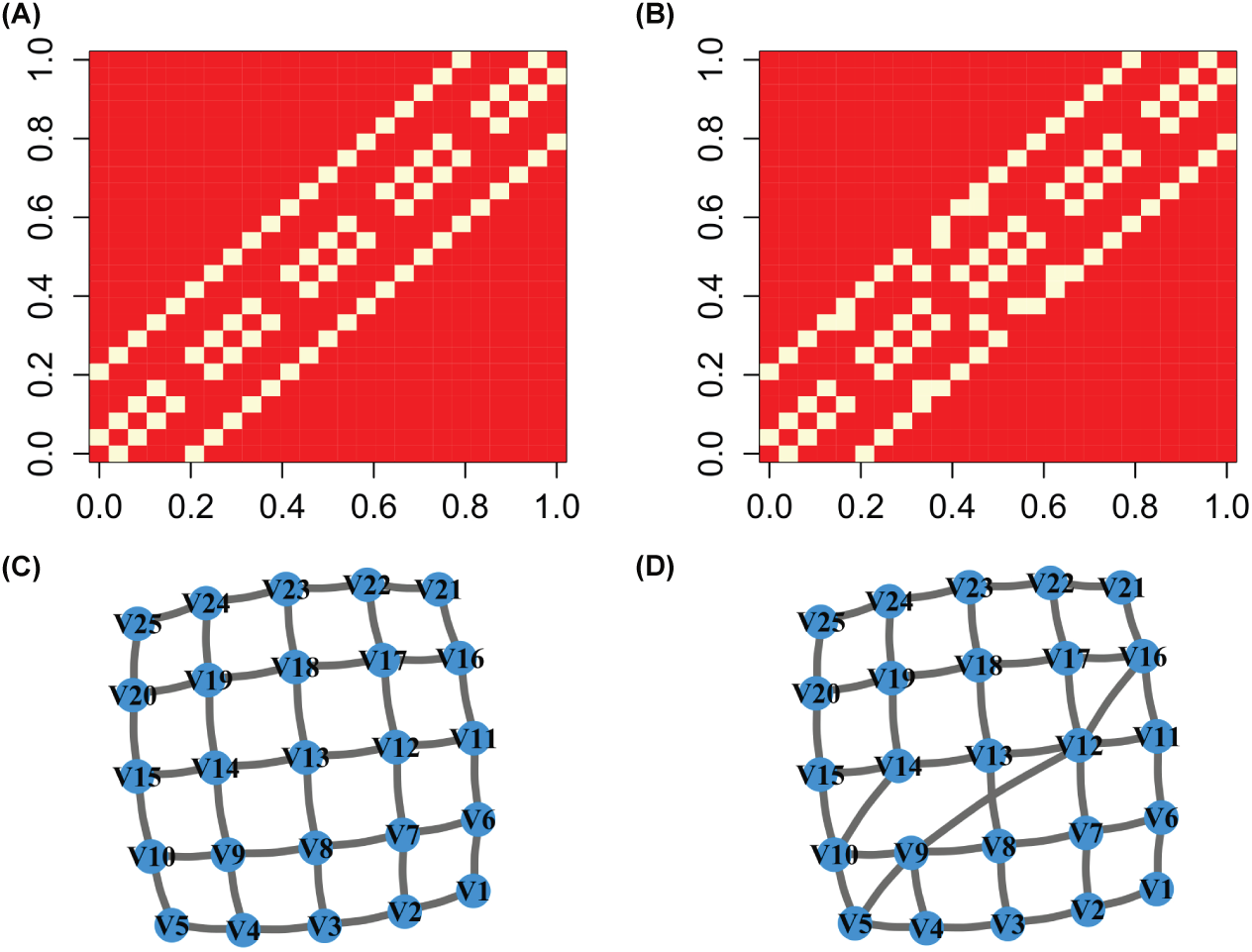
Results of fitting an Ising model to simulated multivariate binary data. The true simulated grid is plotted in (A) and (C). The estimated graph structure via `XMRF(…,method=“ISM”)` is plotted in (B) and (D).

#### Estimate LUSC Mutation Networks

In this section, we estimate the relationships among mutated genes in 178 lung squamous cell carcinoma (LUSC) patients from the TCGA project [16]. We obtained the data via `getTCGA` from `TCGA2STAT` package. A total of 13655 genes for LUSC patients were obtained. Genes with a mutation rate of less than 15% in the cohort or with an undefined gene name were filtered out before analysis. This left data with 59 genes and 179 patients. Similar to the work-flow applied on simulated data presented, Ising models were fit across 20 regularization values, and the optimal network was selected from 100 iterations via the StARS approach.

~~~
> library(TCGA2STAT)
> lusc.mut <- getTCGA(disease=“LUSC”, data.type=“Mutation”)
> mut.dat <- lusc.mut$dat
> mut.rate <- apply(mut.dat, 1, sum)/ncol(mut.dat)
> mut.gd <- mut.dat[mut.rate>= 0.15,]
> mut.gd <- mut.gd[-grep(“Unknown”, rownames(mut.gd)),]
> lusc.mut.fit <- XMRF(mut.gd, method=“ISM”, N=100, nlams=20,
                                    stability=“STAR”, th=0.001, beta=0.1)
> plotGML(lusc.mut.fit, fn=“lusc.gml”, vars=rownames(mut.gd), weight=TRUE)
~~~

The estimated mutated gene networks for lung squamous cell carcinoma viewed from Cytoscape is shown in Figure 6.

**Figure 6.**
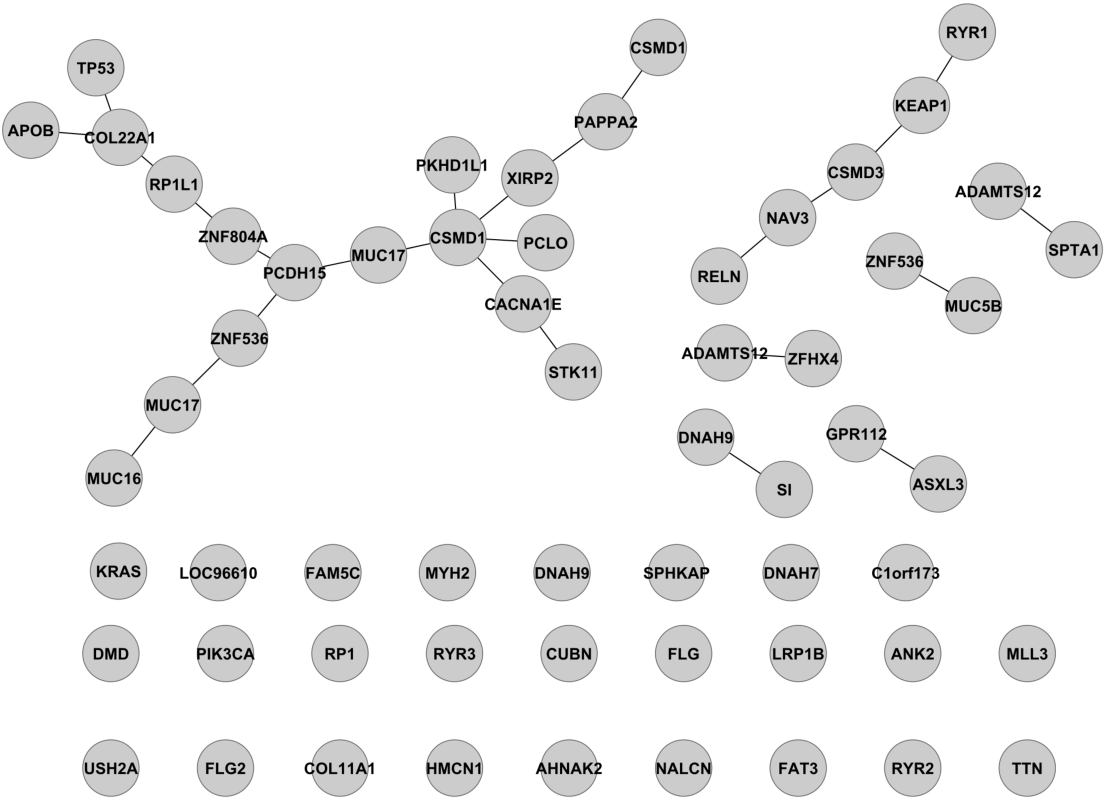
LUSC mutated gene networks estimated by Ising model’s `XMRF(…,method=“ISM”)`

### Tuning Parameter Selection: Network Sparsity

Our `XMRF` package implements two data-driven methods to determine the sparsity of a fitted network. The first method is the stability selection over many bootstrap samples for a single regularization value [7]. The second method is the StARS selection, which is computed over a range of regularization values to select a network with the smallest regularization value that is simultaneously sparse and reproducible in random samples [6]. In this section, both of these methods are demonstrated for an example using the local Poisson graphical model (LPGM).

### Stability Selection

Here, we demonstrate the stability selection technique to learn the network sparsity for networks estimated via the `LPGM` method.

We simulate a scale-free network with 30 variables and 200 observations. We determine the network sparsity based on the stability score, which retains network edges that are estimated in more than 95% (`sth=0.95`) of the 50 bootstrap repetitions (`N=50`). The code is given below:

~~~
> library(XMRF)
> n = 200
> p = 30
# Simulate a scale-free network of 30 notes and 200 samples
> sim <- XMRF.Sim(n=n, p=p, model=“LPGM”, graph.type=“scale-free”)
> simDat <- sim$X
# Compute the optimal lambda
> lmax = lambdaMax(t(simDat))
> lambda = 0.01* sqrt(log(p)/n) * lmax
# Run Local Poisson Graphical Model (LPGM)
> lpgm.fit <- XMRF(simDat, method=“LPGM”, lambda.path=lambda,
                     sth=0.95, N=50)
~~~

Results for the code above are shown in Figure 7. It shows that the estimated network structure (Figure 7(B)) is equivalent to the true network structure (Figure 7(A)). Note that stability selection is the default way to determine network sparsity in `XMRF`.

**Figure 7.**
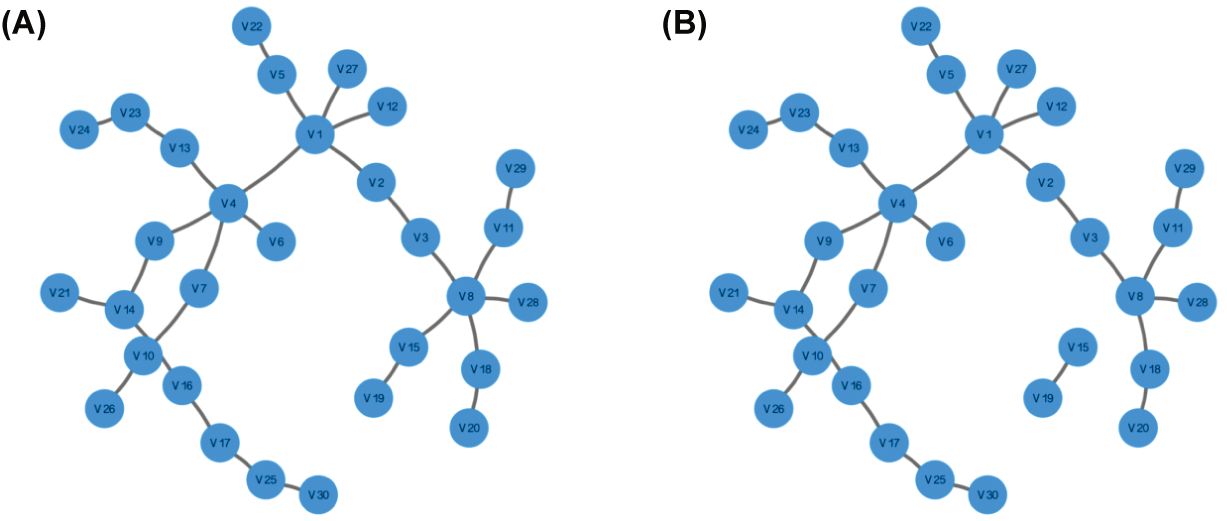
Simulated network from XMRF.Sim(…,model=“LPGM”) (A) and inferred network estimated via XMRF(…,method=“LPGM”) with network sparsity determined via stability selection (B).

### StARS Selection

If users want to fit a network over a series of regularization parameters instead of a single `lambda` as shown in last section, a numerical vector of regularization values should be given for the `lambda.path` parameter of the `XMRF` function.

Another option to study the Markov Networks over the complete regularization path is to let our `XMRF` method decide the path from a null model (empty network) to the full model (saturated network). In this case, the `XMRF(…,method=“”,…)` function will compute the maximum `lambda` that gives the null model and the minimum `lambda` that gives the full model for each of the parametric familes employed. The maximum `lambda` is computed based on the input data matrix, and is the maximum element from column-wise multiplication of data matrix (data matrix in *n × p*) normalized by the number of observations. Based on the maximum `lambda` value, the number of `lambda` (`nlams`) and the minimum `lambda` (`lmin`), sequence of appropriate `lambda` values will be computed.

Stability selection via StARS seeks to select the `lambda` value out of the regularization path which yields the most stable network (or, least variable to bootstrap perturbations). Specifically, the variability of each fitted network is measured based on the stability of edges inferred from the bootstrap samples. The network with the smallest penalization and variability below the user specified cutoff (`beta`) is selected as the final optimal network.

In the following example, we fit the `XMRF(…,method=“LPGM”)` to learn the same simulated scale-free network of 30 nodes from 200 observations along a path of 20 regularization parameters.

~~~
> library(XMRF)
> n = 200
> p = 30
# Simulate a scale-free network of 50 notes and 300 samples
> sim <- XMRF.Sim(n=n, p=p, model=“LPGM”, graph.type=“scale-free”)
> simDat <- sim$X
# Run LPGM on a whole regularization path
lpgm.fit <- XMRF(simDat, method=“LPGM”, nlams=20, stability=“STAR”, th=0.001)
~~~

### Visualization and Data Exportation

To enable users to visualize the inferred network in graphical form, `XMRF` includes three plotting functions with slight variations to serve different purposes. First, the default `plot` function of the `GMS` class will draw the optimal inferred network and save it to a PDF file with the following command:

~~~
> plot(lpgm.fit, fn=“lpgm.fit.net.pdf”)
~~~

Second, the `plotNet` function allows users to plot a specific network with specific layout. For example, to plot the simulated network and the inferred network in Figure 7 with the same layout, the following commands can be used:

~~~
> ml = plotNet(sim$B)
> ml = plot(lpgm.fit, mylayout=ml)
~~~

The third plot function allows users to view the inferred network in other graph visualizing software such as the Cytoscape. The `plotGML` function will write the network in the graph modeling language (GML) format which then can be imported to Cytoscape. For example, with the following command:

~~~
> plotGML(brca.lpgm, fn=“brca.dnet.gml”, weight=TRUE, vars=rownames(brca))
~~~

## Conclusion

We have developed an open source R package that allows users to learn the network structure from data acquired from various high-throughput genomics technologies. Our tool is the only software that allows data to be modeled using their native distribution instead of normalizing the data to follow Gaussian distribution as most other statistical models require. In addition, the parallelization of our algorithms provides an efficient tool for computing large-scale networks.

## Author’s contributions

YW and ZL implemented the R package. GA, PR, EY, and YB developed the theoretical foundation for XMRF.

YW, ZL, and GA wrote the manuscript.

## Acknowledgements

ZL and YW are partially supported by the Houston Endowment and NSF DMS-1263932; GA and YB by NSF DMS-1264058 and DMS-1209017; PR and EY by ARO W911NF-12-1-0390, NSF IIS-1149803, IIS-1320894, IIS-1447574, and DMS-1264033.

